# SOMSC: Self-Organization-Map for High-Dimensional Single-Cell Data of Cellular States and Their Transitions

**DOI:** 10.1101/124735

**Authors:** Tao Peng, Qing Nie

## Abstract

Measurements of gene expression levels for multiple genes in single cells provide a powerful approach to study heterogeneity of cell populations and cellular plasticity. While the expression levels of multiple genes in each cell are available in such data, the potential connections among the cells (e.g. the lineage relationship) are not directly evident from the measurement. Classifying cellular states and identifying transitions among those states are challenging due to many factors, including the small number of cells versus the large number of genes collected in the data. In this paper we adapt a classical self-organizing-map approach to single-cell gene expression data, such as those based on qPCR and RNA-seq. In this method (SOMSC), a cellular state map (CSM) is derived and employed to identify cellular states inherited in a population of measured single cells. Cells located in the same basin of the CSM are considered as in one cellular state while barriers between the basins provide information on transitions among the cellular states. Consequently, paths of cellular state transitions (e.g. differentiation) and a temporal ordering of the measured single cells are obtained. Applied to a set of synthetic data, two single-cell qPCR data sets and two single-cell RNA-seq data sets for a simulated model of cell differentiation, and systems on the early embryo development, haematopoietic cell lineages, human preimplanation embryo development, and human skeletal muscle myoblasts differentiation, the SOMSC shows good capabilities in identifying cellular states and their transitions in the high-dimensional single-cell data. This approach will have broad applications in studying cell lineages and cellular fate specification.

## Introduction

Heterogeneity of cell populations is considered functionally and clinically significant in normal and diseased tissues, and transitions among different subpopulations of cells, such as differentiation, play critical roles during development and disease recurrence [1-3]. In recent years, single-cell gene expression profiling technologies are emerging as increasingly important tools in dissecting heterogeneity and plasticity of cell populations in addition to analyzing cell-to-cell variability on a genomic scale [4]. For example, mammalian pre-implantation development was analyzed from oocyte stage to morula stage in both human and mouse using single-cell RNA sequencing to identify stage-specific transcriptomic dynamics [5,6]; in breast cancer, gene expression profiles of tumor subpopulations along a spectrum from low metastatic burden to high metastatic burden were obtained using qPCR at the single-cell level [7]; and multiple new phenotypes in healthy and leukemic blood cells were defined using gene expression signatures through analysis of single-cell data [8].

Distinguishing or clustering measured cells computationally through their transcriptomic data (e.g. gene expression) is challenging. The number of cells collected in experiments with successful outputs is usually small whereas the number of genes measured usually is significantly larger [9]. In addition, a group of cells collected at one temporal point from one sample may not be perfectly ordered in time compared to the cells collected at slightly different temporal stages, due to cell-to-cell variability in sampling and its nature of unsynchronized cell divisions [10,11]. As a result, a pseudo-temporal ordering of single cells in a high-dimensional gene expression space was introduced [12]. The difficulty in analyzing single-cell data becomes particularly evident for systems of differentiation in which new cell types emerge as time advances, such as the cases of lineage progression during development of murine lung [13] and the differentiation trajectory of skeletal muscles [14].

Ordering single cells temporally, grouping cells of similar transcriptomic profiles, finding transition points, and determining branches are among the key steps in analyzing single-cell data. Clustering methods based on Principle Component Analysis (PCA) or Independent Components Analysis (ICA), such as MONOCLE algorithm [14], group cells according to their specific properties of interests. Several other clustering-based methods such as SPADE [15], t-SNE [16], and viSNE [17] were introduced to identify subpopulations within measured cells without an explicit temporal ordering of the cells. In the Wanderlust algorithm [18], a pseudo-temporal ordering technique incorporated the continuity concept in branching processes, however, with an assumption that cells consist of only one branch during differentiation. To address potential nonlinearity of branching processes in differentiation, a diffusion map technique was adapted to single-cell data by adjusting kernel width and inclusion of uncertainties, enabling a pseudo-temporal ordering of single cells in a high-dimensional gene expression space [19]. With a focus on modeling dynamic changes associated with cell differentiation, a bifurcation analysis method (SCUBA) was developed to extract lineage relationships [20].

Meanwhile, a Waddington landscape of gene expression has been widely used to provide a global and physical view in understanding stem cells and cell lineages [21]. In constructing such landscape, a forward stochastic modeling approach is usually applied to a small gene network with an "energy" function computed through probability density functions or stochastic samplings [22-26]. In this approach, the prior knowledge of the gene regulatory network needs to be known and the landscape is calculated without dimension reduction in the gene space. However, due to computational cost associated with sampling solutions of stochastic differential equations or solving equations of probability density functions of the gene states, the size of network in the landscape calculation usually is small [27].

Here, we propose a new method to analyze single-cell gene expression data by combining a learning method in an artificial neural network (ANN) and a concept similar to a landscape of gene expression data. In this approach, high dimensions of single-cell data are first reduced to two dimensions through a classical unsupervised learning ANN method: the self-organization map (SOM) [28] in which the topological properties of the input data are preserved through a neighborhood function. A cellular state map (CSM) is then derived to mimic a landscape of gene expression data based on a U-matrix calculated by the SOM.

The CSM consists of basins of attractions, which correspond to cellular states, and barriers that separate the different states to indicate directions of transitions between cellular states. Transition paths among the cellular states naturally lead to a pseudo-temporal ordering of the cells. To study effectiveness and capabilities of the method, we apply the self-organization-map for single-cell data (SOMSC) to a set of simulated data and four experimental data sets based on qPCR or RNAseq collected for systems of cell lineages or differentiation.

## Methods

### Preprocess the data

Single-cell gene expression levels measured by qPCR or RNAseq are prone to having missing values, causing bias in analysis without any preprocessing [29]. In this study, we first remove samples that have many zero values in gene expression data. Specifically, the samples of more than 10% of the total number of genes with missing values will not be used; then the missing values of genes in the rest samples are set to the mean value of that gene at its corresponding stage. Another important step in preprocessing is to normalize the data. Because the SOM algorithm uses the Euclidian distance between gene expression vectors of two samples [30], two genes with drastically different ranges of expression values (e.g. expression values of one gene in [0,100] whereas the ones of another gene in the range of [0,0.1]) may influence the SOM unfaithfully, as the larger component may dominate the calculation, introducing bias in analysis. Next we normalize the data linearly such that the variance of each gene is equal to one [30]. The normalized data is stored in a matrix in which each row represents expression values of all genes in one single cell, and the number of rows corresponds to the number of single cells in the data after the preprocessing (Figure 1A).

**Figure 1.**
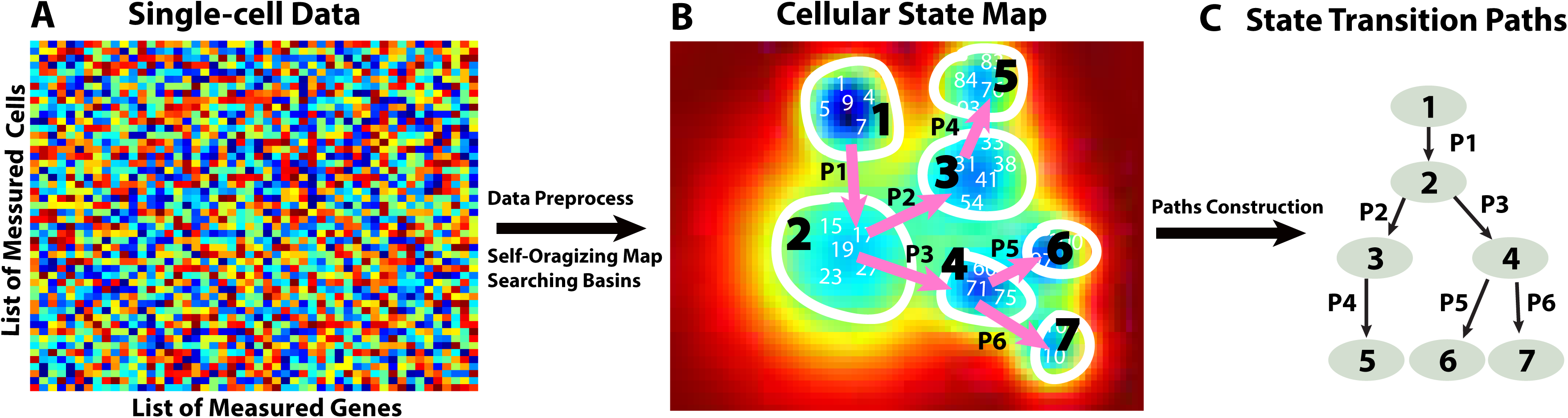
A schematic diagram on constructing cellular state maps (CSMs) and transition paths using the SOMSC method. (A) The gene expression data of single cells. (B) A CSM is constructed by the SOMSC using the data. In the CSM each cell is indexed by a number based on a particular given order or a temporal stage at which the data are collected in measurements. A basin of an attraction in the CSM corresponds to one cellular type. The transitions among different cellular states are labeled by arrows such as P1, P2, ⋯, and P5. (C) The cellular state lineage trees or differentiation processes are then summarized based on the transition path arrows in the CSM.

### Calculate the U-matrix using the Self-Organizing Map

A Self-Organizing Map (SOM) is an effective way of analyzing topology of high-dimensional data, and it projects the data to a low-dimensional surface through a rectangular, a cylinder, or a toroid map [28]. In the SOM, regression of an ordered set of model vectors *m*_*i*_ ∈ ℜ^*n*^ is made into the space of observation vectors *x* ∈ ℜ^*n*^ through the following processes:

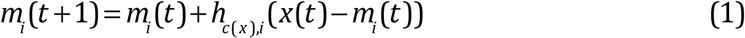
where *t* is an index for a regression step. A regression procedure is performed recursively for each sample *x*(*t*). The scalar multiplier *h*_*c*(*x*),*i*_ is a neighborhood function, acting like a smoothing or blurring kernel over computational grids in the SOM, and often takes a form of Gaussian:

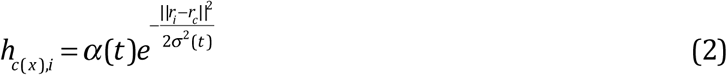
where 0<*α*(*t*)<1 is a learning-rate factor, which decreases monotonically through regression steps; *r*_*i*_ ∈ ℜ^2^ and *r*_*c*_ ∈ ℜ^2^ are locations in the computational grids, and σ(*t*) corresponds to the width of the neighborhood function that also decreases monotonically in each regression step. The subscript *c* = *c*(*x*) is obtained when the following condition is achieved:

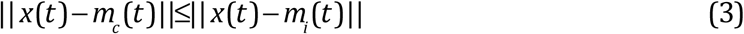

Consequently, *m_c_*(*t*) is the "winner" which matches the best with *x*(*t*). The comparison metric ‖•‖ is selected as the Euclidean metric in Eq.2, and Eq.3. If there are multiple *c*(*t*) satisfying Eq.3 with discrete-valued variables, *m_c_*(*t*) is selected at random for the winner. In the method, a toroid map is used in order to reduce edge effects of the data on the overall mapping [31]. Applying the SOM to the normalized single-cell gene expression data leads to a unified distance matrix (U-matrix) *U*, representing distances between neighboring map units [28].

### Trace the lineage trajectory

#### Construct Cellular State Map (CSM)

To investigate structure of high-dimensional gene expression data, we first define a cellular state map (CSM) *M_cs_* based on the U-matrix *U* through the equation:

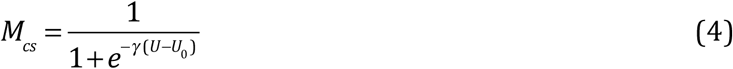

This logistic function transforms *U*, whose elements are always positive, to a matrix *M*_*cs*_, whose elements have values between zero and one. The value of scaling parameter *γ* controls steepness of a sigmoidal curve and the midpoint *U*_0_ determines where 0.5 takes place in the map in Eq.4. The map *M_cs_* may be considered as a Waddington landscape of the high-dimensional gene expression data projected into a two-dimension plane. The basins of attractions of the CSM correspond to individual cellular states in the data.

#### Identify basins of cellular state map

In this process of identifying the basins of the CSM, all local minima in *M_cs_* are searched first, leading to a pool of the minima in an increasing order. To construct the basin of the smallest local minimum (*W*), we first find the smallest local maximum, whose value is denoted as *W_m_*, around this local minimum (*W*). Next we construct contours in the CSM that contains this minimum. The largest such contour value that is still smaller than *W_m_* is the contour that contain the basin of this smallest local minimum (*W*). This searching procedure is then repeated for the second smallest minimum, and the rest of other minima. (More details can be found in Section I in the Supplementary file).

#### Identify transition paths

Cellular state transition paths from one cellular state to the other are traced based on the CSM (*M_cs_*). All cells in the first stage during transition processes need to be known in advance, which is the case for many temporal data. After locating the basins in the *M*_cs_, for the cellular states at the first stage, we then identify its adjacent basins. The neighboring basin that has the smallest height of the barrier is locations of the cells for the next transition state, and then here it means the cells in the basin are at the second stage. If more than one of barriers have the similar heights, indicating a branch process takes place during transitions from the first stage to the second stage, we consider multiple cellular states emerge at the second stage. The procedure consisting of searching for adjacent basins, estimating heights of barriers, and identifying branching processes for each basin continues until all basins are analyzed. At the end of this procedure, the transition paths are also identified (Figure 1BC).

#### Key parameters in SOMSC

In the standard SOM, a two-dimensional U-matrix may have the same size or different sizes in those two dimensions. To avoid bias on a particular gene or a subgroup of genes when applying the SOM to the single-cell data, here we consider both dimensions of a U-matrix to be the same. The total number of grid points in the CSM corresponding to the U-matrix is defined as *N_g_* = *N_r_* × *N_r_* where *N*_*r*_ is the number of grids in each dimension of the CSM. The choice of *N*_*g*_ depends on the number of samples (e.g. the number of single cells), *N*, in order to compute the U-matrix more accurately. Naturally, the size of a U-matrix is proportional to the number of samples, such as *N_g_* = *βN*, where *β* is a constant. Secondly, in the simulation *N_g_* needs to be adjusted to avoid producing too many basins in a CSM, such as the case in which every one or two cells grouped as one basin. Two other key parameters are *γ* and *U*_0_ in a CSM. As shown in the later sections, a CSM seems to produce the most consistent results when the choices of these two parameters enable a larger range of values of elements in *M_cs_* from zero to one, allowing better separation between basins of cellular states.

#### Generate the simulation data

In order to effectively evaluate performance and choices of parameters of the SOMSC, we next construct a toy system consisting of a small number of genes to mimic single-cell gene expression data. There are three stages in the system, and in each stage one type of cells makes a transition to two other types of cells (Figure 2A). Together, seven types of cells with three branches present in the system. The cellular types are defined by the specific patterns of expression levels of the six genes (Figure 2A). Specifically, in Type 1 cells Gene A and Gene B are activated and all other four genes are silenced; in Type 2 cells Gene A, Gene C, and Gene D are activated; in Type 3 cells Gene B, Gene E, and Gene F are activated; when one of Gene A and Gene B and one of Gene C, Gene D, Gene E and Gene F are activated, four other types of cells in the third stage are then defined as Type 4, Type 5, Type 6, and Type 7 cells, respectively.

**Figure 2.**
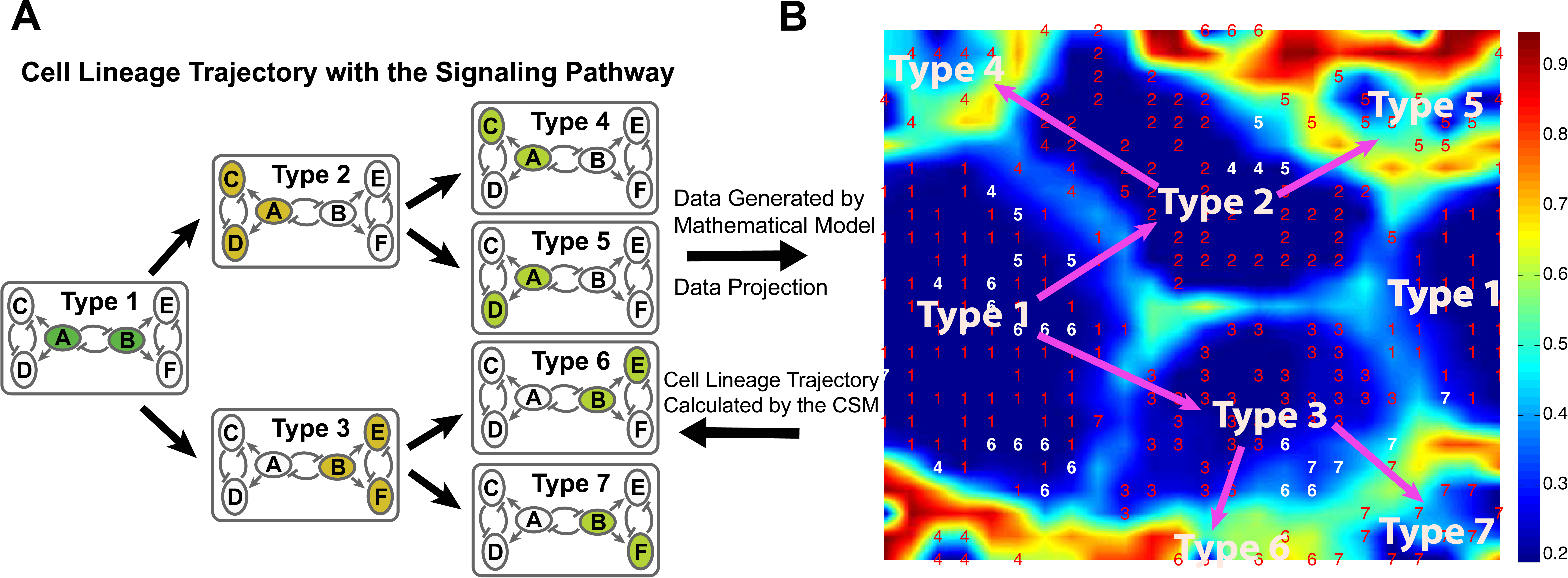
The CSM and cellular state transition paths based on the simulated model. (A) A three-stage lineage system. Stage 1 contains one type of cells in which the activated genes, A and B are highlighted in green; Stage 2 contains Type 2 cells and Type 3 cells. The activated genes, A, C, and D are highlighted in orange in Type 2 cells while the activated genes, B, E, and F are highlighted in orange in Type 3 cells. Stage 4 contains four types of cells: Type 4 cells, Type 5 cells, Type 6 cells, and Type 7 cells. The activated genes, A and C, A and D, B and E, or B and F are highlighted in light green in Type 4, Type 5, Type 6, and Type 7 cells, respectively. (B) The CSM with *N_g_* = 576(24×24) grids is computed for the data of *N* = 353 single cells using *U*_0_= 1.5 and *γ* = 1. A red or white number shown in the CSM is a temporal stage of its corresponding cell in the data. A white number means its corresponding cell locates in an incorrect basin. A pink arrow shows a direction of a transition path.

The system of three-toggle modules consisting of six genes is modeled through a system of stochastic differential equations [19,32,33]. Starting with only Type 1 cells in the system (i.e. the initial state), the expression values of each gene are then collected at three different temporal stages for each stochastic simulation: the early, the middle, and the final stage, in order to mimic a typical set of temporal single-cell data (See Section II in the Supplementary file). Repeating the stochastic simulations using the same set of parameters and the same initial values of genes for 400 times produces a set of gene expression values, corresponding to 1200 sets of single-cell data.

## RESULTS

### SOMSC on the simulation data

To mimic a typical size of experimental data, we randomly select expression levels of 353 cells out of the ones of 1200 cells collected in the simulation data. In the CSM calculated using the SOMSC, each cell is marked by its temporal state collected (Figure 2B). By tracking basins and analyzing heights of barriers, we obtain different cell types and their transition relationship (Figure 2B). Interestingly, in this case the adjacent basins of the basin of Type 1 cells contain all other types of cells from Type 2 to Type 7. However, the barriers between the basin of Type 1 cells and the basins of Type 4, 5, 6, and 7 cells are higher than those for the basins of Type 2 and Type 3 cells, suggesting two possible transition paths: one transition from Type 1 cells to Type 2 cells and the other from Type 1 cells to Type 3 cells (Figure 2B). Next, the barriers between the basin of Type 2 and those of Type 4 and Type 5 are found to be lower than the ones for basins of Type 6 and Type 7 cells. So Type 2 cells make a transition to Type 4 cells or Type 5 cells. The barriers between basins of Type 3 cells and those of Type 6 and Type 7 cells have similar heights, indicating the next transition state of Type 3 cells is either Type 6 cells or Type 7 cells.

To study effects of the number of grids *N_g_* on performance of the SOMSC, we systematically vary *N_g_* and the number of observations *N* in the toy model (See Figure S1). First we fix *N* = 100 observations (or cells) from the toy model but explore five different *N_g_* (See Figure S1A to S1E). When *N_g_* is too small (See Figure S1AB) the CSM is unable to capture all the basins in the system whereas when *N_g_* is too large (See Figure S1E) the CSM tends to overpopulate the basins by grouping every one or two cells into one basin. It is found that the CSM profile becomes more consistent and reliable when *N_g_* is in its middle range of values (See Figure S1C and S1D). Such trend remains when the number of observations (or cells) increases to *N* = 200 (See Figure S1F to S1I), and to *N* = 353 (See Figure S1K to S1O). Together, when *β*, the ratio between *N_g_* over *N*, is in a range of [1,10], the patterns of basins and transition paths in the CSM start to become more consistent. In other words, given the number of observations, the size of the map in the SOMSC *N_g_* needs to be explored until a "convergent" pattern is observed.

It is observed that around 5% of the 353 cells are placed in the incorrect basins in the CSM (marked in white in Figure 2B). Such inconsistency might be due to noise in the data or choices of parameters in the SOMSC. Interestingly, if the data set is analyzed without involving the gene expression levels of those incorrect cells, the new CSM has no cells locating incorrect basins (See Figure S2 and S3 in the Supplementary file), suggesting that either those cells are less consistent compared to the rest of cells in the original data set or the SOMSC is too sensitive to the gene expression levels of those cells.

Two other important parameters in determining the CSM are the midpoint of the logistic function (i.e. *U*_0_) and the scaling factor (i.e. *γ*) in Eq.4. We systematically explore different values of those two parameters and their effects on the CSMs and the transition paths. The sigmoid's midpoint *U*_0_ determines the range of the values of elements in *M_cs_*. A larger value of *U*_0_ usually leads to smaller values of elements of *M*_*cs*_ (e.g. most of elements in *M*_*cs*_ become smaller than 0.5 and some of them are close to zero) while a smaller value of *U*_0_ leads to larger values of elements in *M_cs_* (e.g. larger than 0.5 and close to one). For the scaling factor, a larger value of *γ* usually makes *M_cs_* better cover the entire range of [0,1], however, sometimes it also makes many elements of *M_cs_* close to 0 or 1. It is found that when the elements of *M_cs_* are more evenly distributed in [0,1] by adjusting the parameters *U*_0_, and *γ*, the computed CSM becomes more consistent and reliable (See Figure S4 and S5 in the Supplementary file).

### SOMSC on experimental data

#### qPCR data of mouse embryo development from zygote to blastocyst

Previously, the expression levels of 48 genes at seven time points were measured using qPCR for mouse early embryonic development from zygote to blastocyst [34]. The raw data of the 429 single cells were normalized cell-wisely by the mean expression levels of two genes: Actb and Gapdh [34].

Two different approaches might be applied to such data set by either using the data at each temporal point individually or lumping the data of all seven stages into one set. For example, applying the SOMSC to the data at the second stage results in a CSM with one cell type (Figure 4A), and using the data point at the sixth stage or the seventh stage results in two cell types (Figure 4B) or three cell types (Figure 4C), respectively. However, such approach is unable to determine potential transition paths among cell types inherited in the data because different basins or cellular states are obtained using different CSMs.

**Figure 3.**
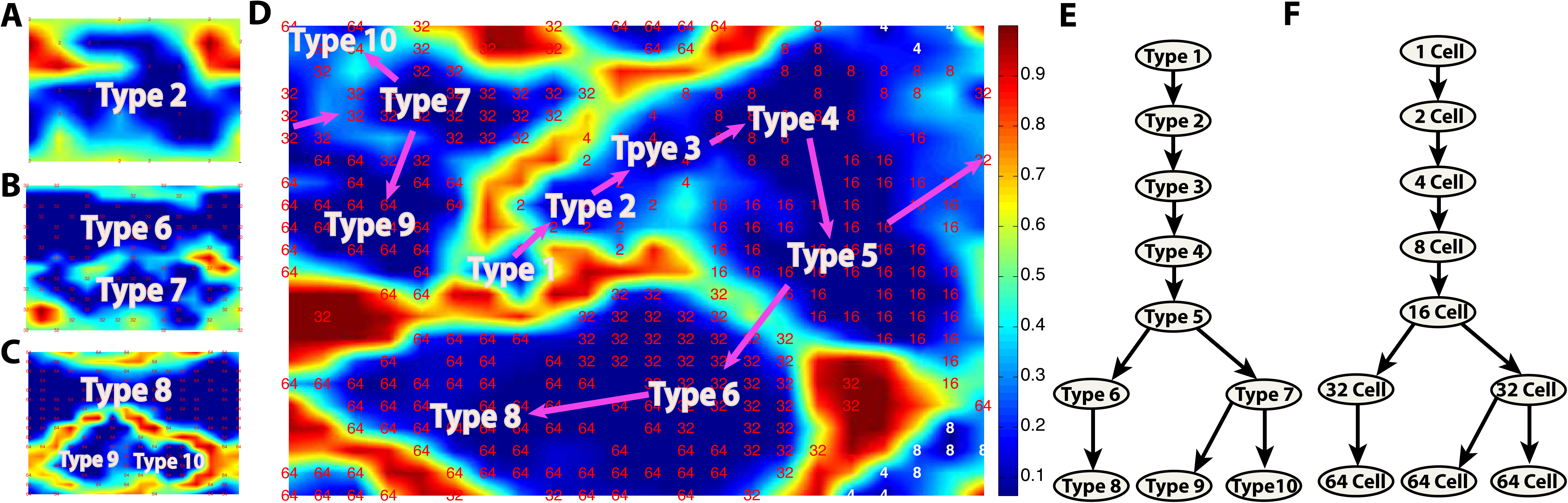
CSMs and a lineage trajectory are constructed using the qPCR data of mouse stem cells from zygote to blastocyst [48]. (A-C) CSMs obtained using data only at the second, sixth and seventh stages, respectively. A red or white number in (A, B, C) represents an index of stages when the expression levels of cells were measured. (A) Type 2 labels the only basin of cells in the CSM computed using the data only from the second stage. Here *N_g_* = 36, *U*_0_= 0.5 and *γ* = 0.01. (B) Type 6 and Type 7 label two separate basins of the CSM computed using the data only from the sixth stage. Here *N_g_* = 196, *U*_0_= 2 and *γ* = 0.3. (C) Type 8, Type 9, and Type 10 label three separate basins of the CSM using the data only from the seventh stage. Here *N_g_* = 196, *U*_0_= *2* and *γ* = 0.3. (D) The CSM is computed using the data collected all seven stages with a total of *N* = 442 cells. Here *N_g_* = 484, *U*_0_= *2* and *γ* = *2*. Ten basins are labeled by Type 1, Type 2, ⋯, and Type 10. A white number means its corresponding cell is located in an incorrect basin. A pink arrow indicates a direction of a transition path. (E) The state transition paths are derived from the CSM in (D). (F) The differentiation lineage tree of early mouse development was obtained in a previous study [48].

**Figure 4.**
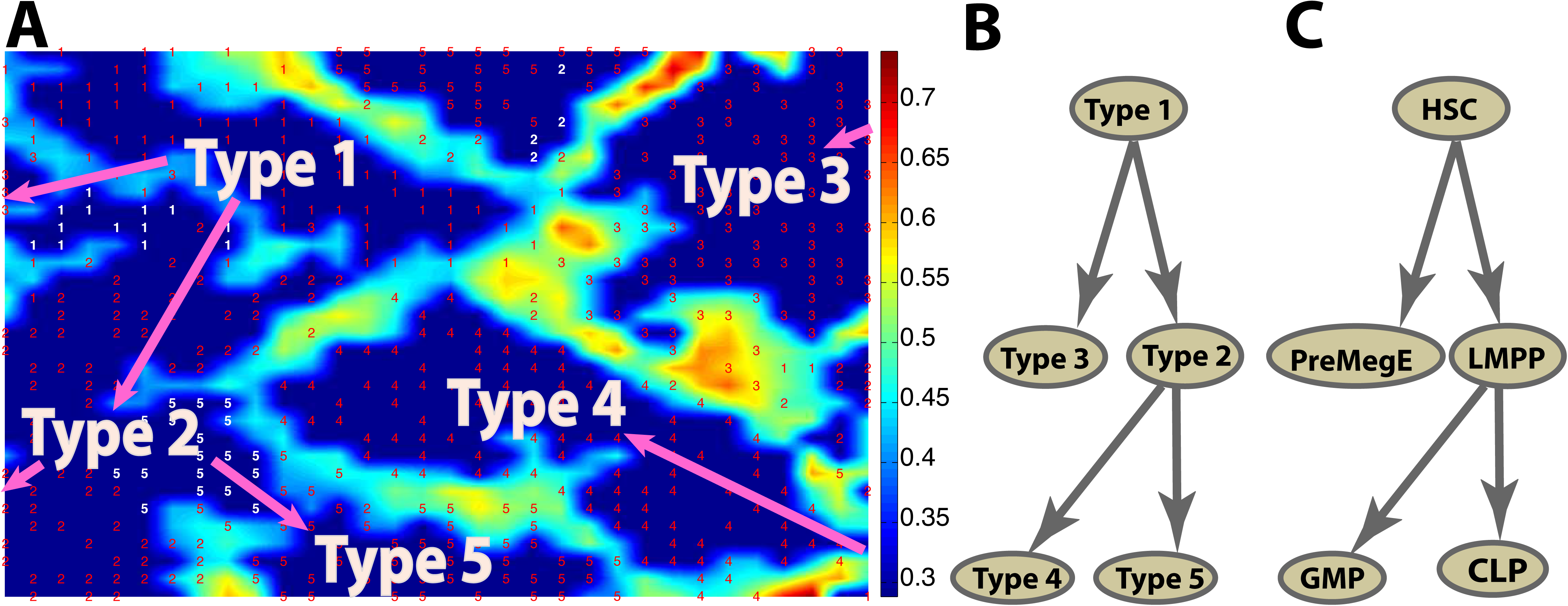
The CSM and a cell lineage trajectory are constructed using the qPCR data of mouse haematopoietic stem cells [35]. (A) The CSM is computed using *N_g_* = 1024 based on *N* = 597 cells. Here *U*_0_= 1.5 and *γ* = 0.88. A red or white number represents a cell with a specified type given in the single-cell measurement [35]. The cells marked in white numbers are those in incorrect basins. A pink arrow is a direction of a transition path. (B) The state transition paths are obtained from the CSM in (A). (C) The lineage tree of mouse haematopoietic stem cells was obtained in the previous study [35].

Using all 442 cells collected at the seven stages simultaneously produces one CSM containing 10 basins (Figure 4D), and the relationship of those basins can then be analyzed to study state transitions. The basin labeled as a Type 1 cell is chosen based on those cells marked at the initial stage in the collected data [34]. The other nine basins are labeled by Type 2, ⋯, Type 10. In the CSM, the Type 1 cell has three neighboring basins, and the barrier between the basin of the Type 1 cell and the basin of the Type 2 cell is found to be lower than those barriers separating with other basins, indicating the Type 1 cell makes a transition to the Type 2 cell. Similar analysis suggests that the Type 3 cell is the next transition state of the Type 2 since the corresponding barrier height is lower than others.

As seen in the CSM, clearly there is a transition from the Type 3 cell to the Type 4 cell. The height of the barrier between the basin of the Type 5 cell and the basin of the Type 4 cell is lower than others, showing that the Type 4 cell makes a transition to the Type 5 cell. The next transition states of the Type 5 cell are the Type 6 cell or the Type 7 cell because the heights of the barriers between them are lower than others, suggesting a branch process takes place. The barrier between the Type 8 cell and the Type 6 cell is rather low, indicating that the Type 6 cell becomes the Type 8 cell. Finally, two basins adjacent to the Type 7 cell have barriers of similar heights, indicating that there are two transitions from the Type 7 cell to the Type 9 cell or the Type 10 cell. As a result, seven stages containing two branches are identified, corresponding to the seven developmental stages [34]: 1-cell stage, 2-cell stage, ⋯, 64-cell stage. Two major cell types (TE and ICM) arise at the 32-cell stage, and later the ICM cells differentiate to EPI or PE cells at the 64-cell stage (Figure 4E). To investigate each individual cell, one can index each cell by a proper order to scrutinize its location in the CSM for its transition capabilities or other properties relative to some other cells (see Figure S6 in the Supplementary file).

It is not surprising that a very small number of cells (around 5% out of 442 cells marked in white color) that were collected at one developmental stage in the experiment are not exactly located in the corresponding basins of the CSM (Figure 4D). Interestingly, the "mismatch" cells are found to be mostly collected in the 8-cell stage. Noise in the measurements, the small number of observations, and the choices of parameters used in the SOMSC may all contribute to this mismatch. To further study this, we next vary the sizes of mappings from *N_g_* = 484 to *N_g_* = 900, and find that the overall patterns of the lineage trees hardly change (See Figure S7 in the Supplementary file). However, when we use *N_g_*= 100 or *N_g_* = 3600, the number of basins and the obtained transition paths start to become inconsistent (See Figure S8 in the Supplementary file). Overall, it is important to vary the parameters used in the SOMSC in order to capture a reliable CSM with consistent cell types and transition paths using the noisy single-cell data.

#### qPCR data of mouse haematopoietic stem cells

In a previous study the expression levels of 24 genes including 18 core transcription factors were measured using qPCR for 597 mouse haematopoietic and progenitor stem cells [35]. The data were then normalized to the mean expression levels of two genes: Ubc and Polr2a [35]. After applying the SOMSC to this data set, we observe five different basins, indicating five possible cellular states inherited in the data marked by Type 1, Type 2, ⋯, Type 5 (Figure 5A). The Type 1 cell is identified using the prior knowledge given in the data [35]. Comparing all barriers surrounding the Type 1 cell, the height of barriers for Type 2 and Type 3 are much lower than the others. However, the height of the barrier for the Type 2 cell and the Type3 cell is similar, suggesting that the Type 1 cell may become either the Type 2 cell or the Type 3 cell. Similarly, it is found that the Type 2 cell may make a transition to either the Type 4 cell or the Type 5 cell.

**Figure 5.**
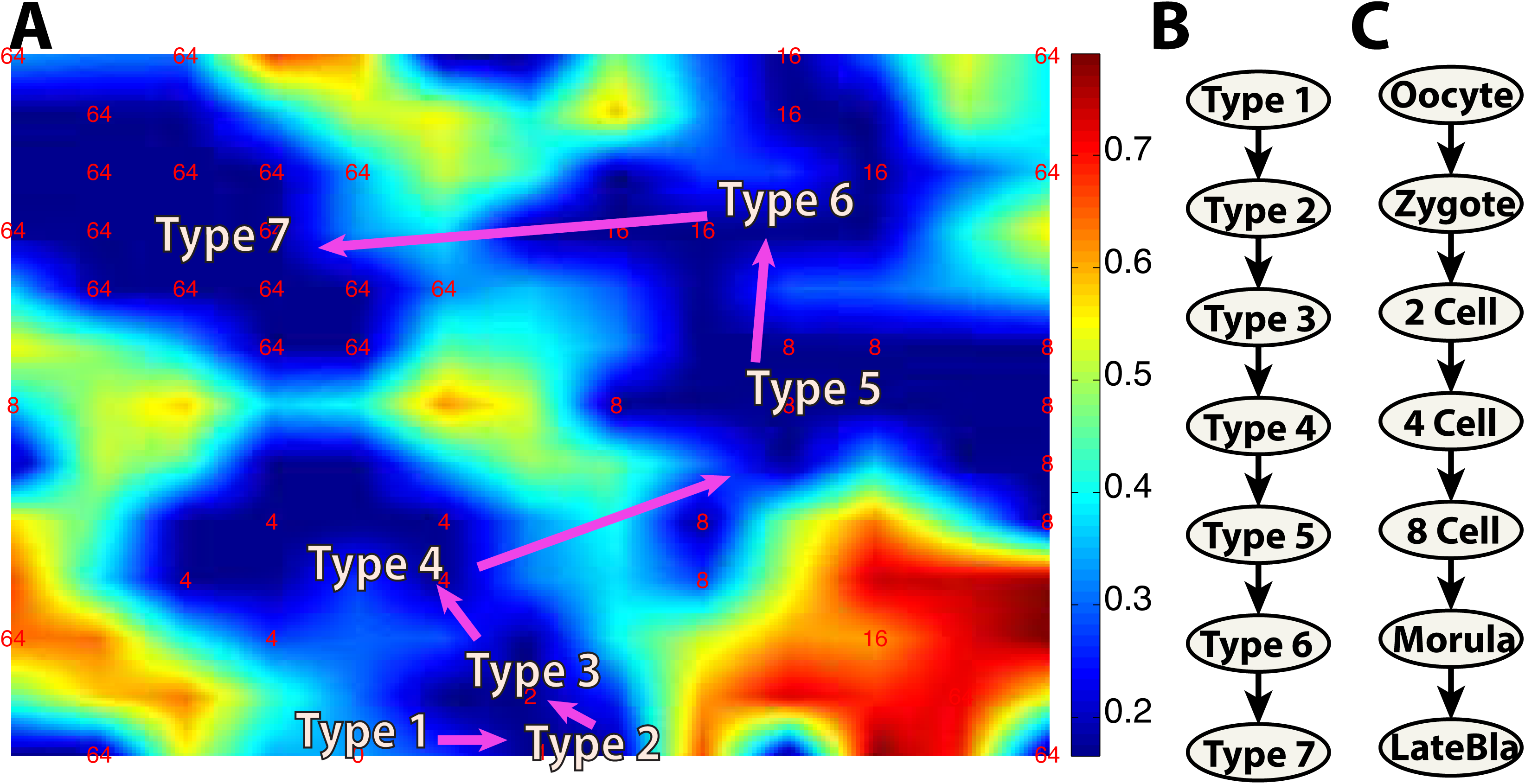
The CSM and lineage relationship are constructed using the RNA-seq data of human preimplantation embryonic cells from oocyte to late blastocyst [6]. (A) The CSM is calculated using *N_g_* = 169 based on all *N* = 90 cells collected at differentiation stages. Here *U*_0_= 20 and *γ* = 0.1. A red or white number represents a stage of a cell measured. A white number means its corresponding cell is located in an incorrect basin. A pink arrow is a direction of a transition path. (B) The paths of transition are calculated from the CSM in (A). (C) The differentiation lineage tree of human preimplantation embryonic cells was obtained in the previous study [49].

Once the transition paths of the five types of cells are obtained (Figure 5B), we can easily establish a map between the transition paths and the well-known lineage trajectory of five mouse haematopoietic cell types [35]: haematopoietic stem cell (HSC), lymphoid-primed multipotent progenitor (LMPP), megakaryocyte-erythroid progenitor (PreMegE), common lymphoid progenitor (CLP) and graulocyte-monocyte progenitor (GMP) (Figure 5C).

Similar to the previous cases, a very small portion of cells fall into the incorrect basins (Figure 5A and Figure S9 in the Supplementary file). For example, a small number of HSC cells (marked by white numbers in Figure 5A) are found located in the basin of the LMPP cells whereas a small number of CLP cells (also labeled in white) are found in the basin of LMPP cells. Missing entries in the raw data, the pre-processing method [35], the fact that LMPP is the intermediate cell types during transitions, and our choices of parameters in the SOMSC may all contribute to the mismatch. Also, similar to the study on the toy model, the choice of proper *N_g_* is important in tracking the transition paths, and too small or too large values of *N_g_* lead to inconsistent patterns of the CSMs (See Figure S10 in the Supplementary file).

#### RNA-seq of human preimplantation embryos

In a previous single-cell RNA-seq analysis on human preimplantation embryos, 90 individual cells were sorted at seven stages: metaphse II oocyte, zygote, 2-cell, 4-cell, 8-cell, morula and late blastocyst, with two or three embryos used at each stage [6]. In this study, over 20,000 genes were measured using RNA-seq. Because the number of cells is small and the number of genes is very large in the data set, we only select those genes that are significantly expressed at least at one stage, leading to a system of 2,389 genes and 90 cells.

A CSM calculated by the SOMSC contains seven basins of cells (Figure 6A). A Type 1 cell is identified based on those cells in the metaphase II oocyte [6]. The rest of basins are then labeled by Type 2, Type 3, ⋯, Type 7. The barrier between the Type 1 cell and the Type 2 cell is found lower than those for the Type 3 cell, and the Type 7 cell. It indicates that the Type 1 cell make a transition to the Type 2 cell. Comparing the heights of barriers among the adjacent basins, the Type 2 cell likely make a transition to the Type 3 cell, and the next transition state of the Type 3 cell is the Type 4 cell that can make a transition to the Type 5 cell. Similar analysis shows that the Type 5 cell becomes the Type 6 cell that makes a transition to the Type 7 cell (Figure 6B). The observed cellular states and transition paths are consistent with the previous study (Figure 6C) [6]. The location of each cell and the distribution of cells in the CSM potentially provide additional information (e.g. signature genes for specific cellular types) for the lineage tree (See Figure S11 in the Supplementary file).

**Figure 6.**
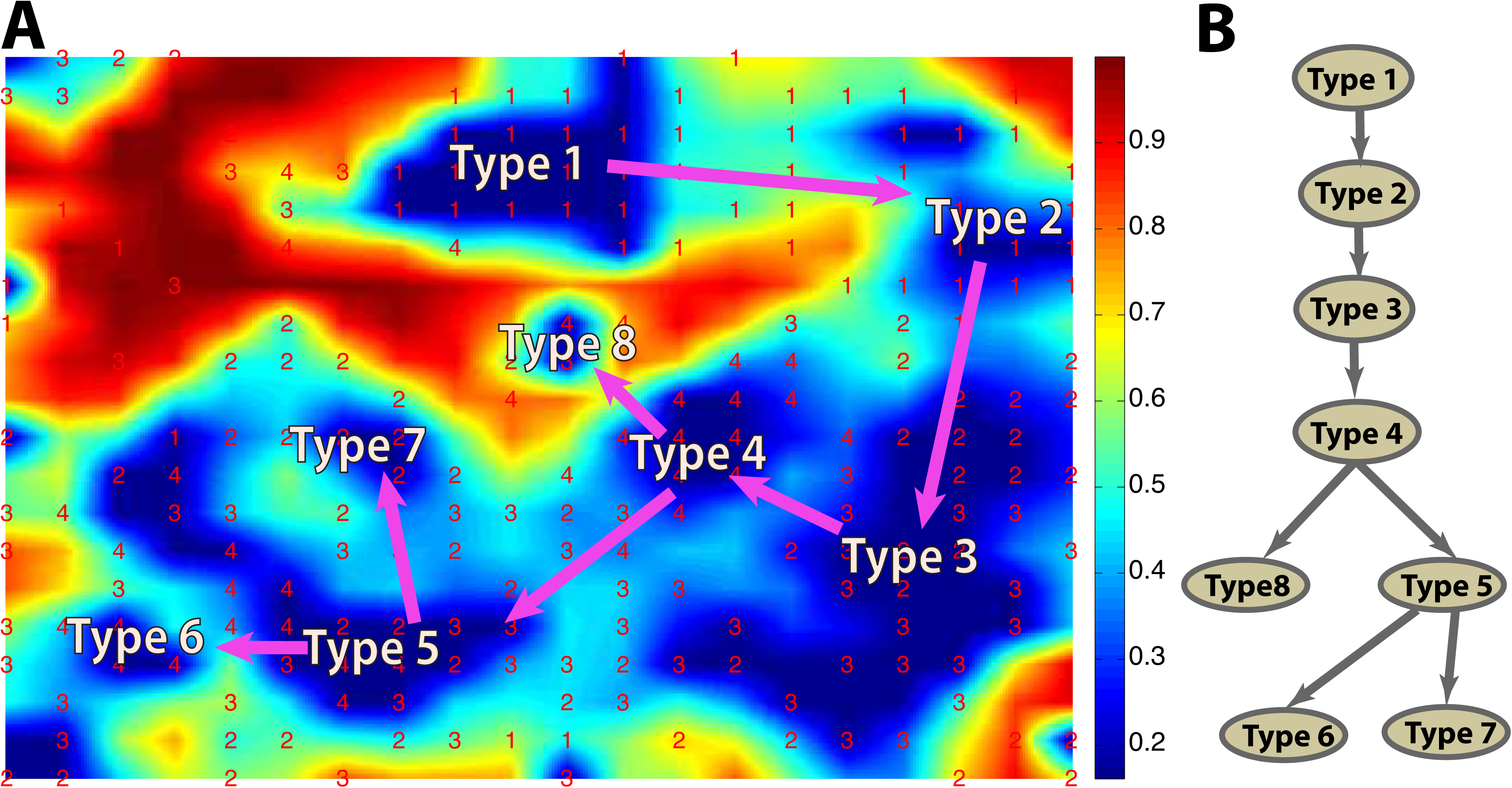
The CSM and lineage transition relationship are constructed using the single-cell RNA-seq data from human skeletal muscle myoblasts [12]. (A) The CSM is calculated using *N_g_* = 400 based on *N* = 271 human skeletal muscle myoblasts cells collected at 0h, 24h, 48h, and 72h. Here *U*_0_= 5 and *γ* = 0.8. A red number is an ordered time point when the expression levels of cells were measured. The pink arrow is the direction of the transition path. (B) The lineage tree is predicted based on the CSM in (A).

It is found that too small or too large *N_g_* in the SOMSC may result in inconsistent patterns of basins and transition paths in the CSMs (See Figure S12 in the Supplementary file). However, by tuning the parameters in a systematic way, the SOMSC is able to obtain a "convergent" CSM and transition patterns.

#### RNA-seq of human skeletal muscle myoblasts

In a previous study single-cell RNA-seq of 271 cells collected from differentiating human skeletal muscle myoblasts (HSMM) were measured at 0, 24, 48 and 72h after switching human myoblasts to low serum [12]. 518 genes that were significantly and differently expressed across different time points and considered to be associated with myoblast differentiation were measured [12].

In the CSM consisting of seven basins marked by Type 1, Type 2, ⋯, Type 8 (Figure 7A), The Type 1 cell and the Type 2 cell were collected at 0h [12]. Analysis on the heights of barriers shows that a transition takes place from the Type 2 cell to the Type 3 cell, which can becomes the Type 4 cell. There are two adjacent basins next to the Type 4 cell, which may make a transition to the Type 8 cell or to the Type 5 cell. Finally, the Type 5 cell can become either the Type 6 cell or the Type 7 cell. The transition paths in a form of a lineage tree are then constructed accordingly (Figure 7B).

By comparing the temporal stage marked on each cell and the cell types identified using the SOMSC, we find that the transitions predicted from Type1, along Type 2, and Type 3, to Type 4 is consistent with the temporal sequence shown in the data. The CSM also predicts two different types of cells at 0h: Type 1 and Type 2, indicating a mixture of two subpopulations of cells at 0h. In addition, Type 3 consists of cells collected at both 24h and 48h. The CSM shows two branching processes taking place from the Type 4 cell to the Type 5 cell or to the Type 8 cell, and from the Type 5 cell to the Type 6 cell or to the Type 7 cell. The two branches are similar to those obtained by other algorithms [12,36]. It is interesting to note that there are four types of cells collected at 24h, three types of cells collected at 48h, and three types of cells collected at 72h. These mixtures of different types of cells in multiple temporal stages suggest the gene expression plasticity might take place between the time points of measurements. Together, our simulations show capabilities of the SOMSC in predicting multiple cellular states and potential plasticity of subpopulations of cells.

## Conclusion and Discussion

In this paper we have presented a self-organization-map based method for analyzing single-cell gene expression data that may contain multiple cellular states with transitions among them. Applications of the SOMSC to a set of simulated data and four sets of differentiation data have demonstrated strong capabilities and effectiveness of the SOMSC in identifying cellular states and their transitions.

A cellular state map (CSM) based on a U-matrix calculated from the SOM provides a global landscape view of cell differentiation or cellular state transitions. By estimating the heights of barriers between basins in a CSM, transition paths among the states are then identified. The location of each cell in the CSM may provide useful information on the cell's viability and potential of transitions to different cellular states. Such knowledge on individual cell in single-cell data is lacking in many other methods for single-cell analysis.

The major computational cost of the SOMSC comes from the iteration procedure in calculating the U-matrix in the SOM, with a complexity of O(*NN_g_DT*) where *D* is the number of genes measured in the data, *T* is the number of iterations used in the SOM, and *N* is the number of samples in a single-cell data set [37]. In practice, *D* is usually around 1,000 (the number of genes significantly expressed), and both *T* and *N* are less than 1,000, implying a complexity of Ο(10^9^) that the SOMSC is able to handle effectively.

Single-cell data are often used to identify cellular states in heterogeneous populations of cells [38]. However, the complexity in data visualization and analysis presents a major difficulty in distinguishing such subpopulations. The SOMSC may capture complex topological shapes in the data to identify those subpopulations due to the advantageous feature of the SOM unlike many other methods requiring convex or normal structure of the data [39]. Another major feature of the SOM is its capability of finding multiple minima as the entire space of feasible solutions in the SOM is searched until finding optimal solutions [39,40]. This is consistent with the observations that the SOMSC is rather stable in searching for basins of attractions and transition paths in the CSM of single-cell data.

Several parameters in the SOMSC need to be tuned in order to obtain a reliable CSM. It is not surprising that given a number of samples (the number of cells and the number of genes measured), the number of grids for a U-matrix calculated by the SOM requires adjustment in order to obtain "convergence" of a corresponding CSM. The scaling parameter *γ* in Eq.4 of a CSM was found to reduce noise effects in a U-matrix, allowing well-separated basins and well-defined barriers. Another important element to improve in the SOMSC is the approach in identifying basins and barriers. Matlab built-in contour construction method is currently used in this paper, and other algorithms may be further explored.

Noise and variability in single-cell data introduce another major complexity. In this work we have tried to reduce noise and variability effects by first removing those identified 'noisy' data from the training data sets. For example, in the case of the simulation data, cells located in incorrect basins are considered as the 'noisy' data. While a similar approach might be used for experimental data, identification of incorrect basins is clearly challenging, depending on availability of appropriate experimental measurements and prior knowledge on the systems. Potentially, machine-learning methods might be explored to enable reduction of noise effects for constructing a more consistent CSM. Other possibilities of improvement in this area include usage of different distance metrics (e.g. the diffusion metric [19]) instead of the standard Euclidean distance metric used in this work.

Previous works demonstrated that the confounding errors (e.g. batch errors) have great effects on single-cell data [29,41]. PCA [42], surrogate variable analyses [43], probabilistic estimation of expression residuals [44,45] or removal of unwanted variation [46] were explored to reduce such effects of confounders in gene expression measurements of the bulk cell populations [47]. Potentially, those methods could be extended to single-cell data. Other factors that are more unique to single-cell measurements, such as cell division, which may induce cell-cell variability, will provide an additional difficulty, for which a linear mixed model could be utilized [29]. In general, reducing the effects of confounding errors is essential to producing reliable classification of cellular states and identifying the transition paths among them.

A CSM produced by the SOMSC is similar to the gene expression landscape although a typical landscape is a function of each gene without dimension reduction. It would be interesting to make a comparison between a landscape computed by forward modeling based on a small size of network and a CSM generated by the SOMSC on single-cell data. Overall, the SOMSC provides a robust and convenient approach to classify the cellular states and to identify their transitions, and it is powerful in suggesting signature transcription factors, branching processes, and pseudo temporal orders of single cells.

